# Positive relationships between interspecific association strength and elevational range overlap highlight the role of facilitation in community assembly

**DOI:** 10.64898/2026.11.06.764909

**Authors:** Sarthak Mondal, Shambu Rai, Bharat Tamang, Aman Biswakarma, Dambar Kumar Pradhan, Mangal Rai, Aiti Thapa, Dema Tamang, Kabir Pradhan, Vikram Rai, Umesh Srinivasan

## Abstract

The interplay of abiotic conditions and biotic interactions determines individual species’ ranges, and by extension therefore, community assembly. To date, studies have focused predominantly on abiotic filtering (e.g., temperature) and agonistic interactions (e.g., competition) as potential drivers of community assembly; the role of facilitation in setting geographical range limits and structuring communities remains unclear. We tested how participation in compositionally distinct modules of facilitative networks of interactions in mixed-species bird flocks (MSFs) influenced interspecific elevational range overlaps across a 2000m elevational gradient in the hyperdiverse Eastern Himalaya. Species that participated in MSFs had higher average elevational range overlap compared with MSF non-participants. Positive correlations between pairwise association strength in MSFs and elevational range overlap were found for all MSF modules. We highlight the potential role of facilitation in determining species co-occurrence patterns and assembling communities across large spatial gradients.

## Introduction

The processes governing the limits of species’ geographical ranges have long been of interest (dating back at least 200 years; Humboldt & Bonpland 1807, Darwin 1859) and understanding such processes remains foundational to ecology. Strong evidence indicates that historical evolutionary processes act in concert with current ecological factors to determine a species’ range (Geber 2011); species-specific physiologies determine species’ fundamental niches (the abiotic conditions under which a species can maintain population growth rates ≥ 1; Hutchinson 1957), whereas biotic interactions such as competition and facilitation shape the realised niche (where the species actually occurs in space and time; Sexton et al. 2009, Pigot & Tobias 2013, Cahill et al. 2014, Harsch & HilleRisLambers 2016). Earliest theory focused on abiotic factors such as temperature to explain species’ range limits (Humboldt & Bonpland 1807, Merriam 1894, Gleason 1926), arguing that a species can persist and maintain a viable population within its geographic range only where environmental conditions are suitable. For instance, at very low and high temperatures, survival and reproduction will be low, whereas performance and population growth rates are highest at intermediate optimal temperatures, thereby setting range limits at the margins of species-specific thermal tolerances (Louthan et al. 2015).

Investigating the role of biotic interactions as determinants of range limits has been comparatively recent, likely because of the inherent difficulty in quantifying the effects of species interactions on participant species, and the complex interaction milieu within which pairwise interactions are embedded (Gaston 2003, Sexton et al. 2009). Further, biotic interactions between species pairs are often highly context-dependent (Callaway et al. 2002, Ohgushi et al. 2012, Schöb et al. 2014, Cai et al. 2020), and uncovering the role of species interactions in setting range limits remains challenging. Historically, studies examining the role of biotic interactions in determining species’ ranges have focused mainly on agonistic interactions such as competition and predation, in which at least one participant in the interaction faces a negative outcome from that interaction (Connell 1983, Bull & Possingham 1995, Case & Taper 2000, Holt et al. 2011). Facilitative interactions such as commensalism and mutualism/symbiosis, in which at least one partner benefits, can also influence range limits (Stephan et al. 2021), but remain poorly understood as determinants of geographic range.

Positive interspecific interactions help species to maintain populations in space and time under conditions that might otherwise be unsuitable by, for instance, enhancing resource acquisition and environmental tolerance, and promoting the persistence of rare species that might otherwise be outcompeted (Catalayud et al. 2020, Stephan et al. 2021). Depending on the intensity of such interactions, positive interactions can shape distributions in myriad ways. – An obligate mutualism between two species (such as a highly specialist plant-pollinator interaction) would be expected to result in nearly complete range overlap between the plant and its pollinator (Pauw & Bond 2011, Moeller et al. 2012, Harrower & Gilbert 2018). On the other hand, facultative beneficial interactions could increase the realized niche of participant species, leading to larger species’ geographic ranges than expected in the absence of such an interaction. Such a phenomenon has been well documented in nurse plants, which help native and non-native plants establish ranges in novel and harsh environments (Brittingham & Walker 2000, Cavieres et al. 2008).

Mixed-species bird flocks (MSFs) are highly cohesive groups of individuals from multiple (insectivorous and omnivorous) species that forage and move together and are widespread in forests worldwide (Sridhar et al. 2009, Goodale et al. 2017). MSFs occur year-round in regions such as the Amazon, but only in winter or dry season in most other parts of the world (Srinivasan et al. 2012, Mangini & Areta 2018, Jones et al. 2020, Rutt & Stoufffer 2021, Mangini et al. 2023, Martínez et al. 2023). Within MSFs, multiple species form complex networks of obligate or facultative facilitative interactions, and participation in MSFs can increase individual fitness (Jullien & Clobert 2000, Srinivasan 2019) through anti-predator and foraging benefits (Waite & Grubb 1988, Martínez et al. 2018). Reduced predation risk can arise from several mechanisms such as dilution (i.e., decreased statistical probability of any one individual to be depredated upon), the confusion effect (difficulty of predator in targetting a specific individual in a group) and the many eyes effect (increased probability of detecting a predator; Pulliam 1973, Foster & Treherne 1981, Terborgh 1990). In addition, MSFs often contain “sentinel species” that excel at predator detection and provide early warning (Munn 1984, Terborgh 1990). Foraging benefits might arise from prey being flushed from the vegetation by gregarious species, making them more accessible to individuals in the MSF, or via the exchange of information regarding suitable foraging locations (Winterbottom 1943, Waite & Grubb 1988, Beauchamp 2005, Sridhar et. al. 2009). Species interactions in MSFs can influence community processes (Goodale et al. 2015, Goodale et al. 2020) and MSFs can therefore serve as a useful model system to investigate the role of facilitative interactions in shaping community-level patterns such as overlaps in geographical range (Clements 1916, 1936). For instance, multi-species territoriality, in which participant species in MSFs jointly defend home ranges against conspecifics in neighbouring MSFs, might arise from strong facilitation between functionally important “nuclear species” and other participants (Munn & Terborgh 1979, Martínez et al. 2018).

In mountains, elevational gradients are relatively simple systems to understand geographical ranges (Maitra et al. 2026) – that they are one-dimensional, and every montane species has a distinct lower and upper elevational range limit. MSFs change predictably in composition (Weske 1972, Jankowski 2013) with elevation, with low-elevation MSFs generally comprising different species than mid-elevation MSFs and in turn, high elevation MSFs, although many species with large elevational ranges can occur in MSFs across elevations (Montaño-Centellas et al. 2021, Montaño-Centellas et al. 2023). In addition, even at any given elevation, species often segrate into distinct MSF types based on traits such as habitat specialisation (Srinivasan et al. 2026), body mass, plumage colour and foraging stratum (Srinivasan et al. 2012), likely because similarly size and colour can reduce predation risk through dilution and confusion (Goodale et al. 2020). We examined how MSF participation was related to species distributions across a large elevational gradient in the hyperdiverse Himalayan mountain range, the tallest on earth. We hypothesised that because of facilitative interactions between species pairs within compositionally different MSFs across an elevational gradient, species that co-occur in MSFs are also likely to co-occur closely in elevation.

Following from this hypothesis, we predicted that:

1. Overlaps in the elevational ranges of species co-participating in MSFs should be significantly higher than range overlaps between species that do not participate in MSFs (and therefore cannot exchange facilitative benefits).
2. The strength of association between pairs of MSF participant species should be positively correlated with the degree of elevational range overlap because greater benefits being exchanged between species should lead to greater co-occurence in MSFs (i.e., species’ association) and space (i.e., range overlap).
3. Elevational range overlap will be higher between species pairs within distinct MSF types/modules (e.g., large-bodied versus small-bodied MSFs) compared with overlap between *all* MSF participating species. Further, the strength of correlation between association strength and range overlap for species pairs within distinct MSF types should be stronger than the correlation for *all* MSF participating species.

## Results

We analysed a total of 996 MSFs (180 MSFs at 800m, 175 at 1200m, 167 at 1600m, 170 at 2000m, 137 at 2400m and 167 at 2800m). In total, the analysis consisted of 84 species that participated in MSFs and 158 non-flocking species, resulting in a combined species pool of 242 species.

### Characterisation of unique MSF modules using cluster analysis

Using cluster analyses, we identified distinct MSF types or “modules” found at each elevation (Fig. 1). Out of these, we found nine unique MSF types that contained 10 or more participant species across the elevational gradient. Amongst these, MSFs at 800m, 2400m and 2800m were found to have one highly speciose MSF type whereas MSFs at 1200m, 1600m and 2000m were each found to have two highly speciose MSF modules (Fig. 1, Table S2).

**Figure 1.**
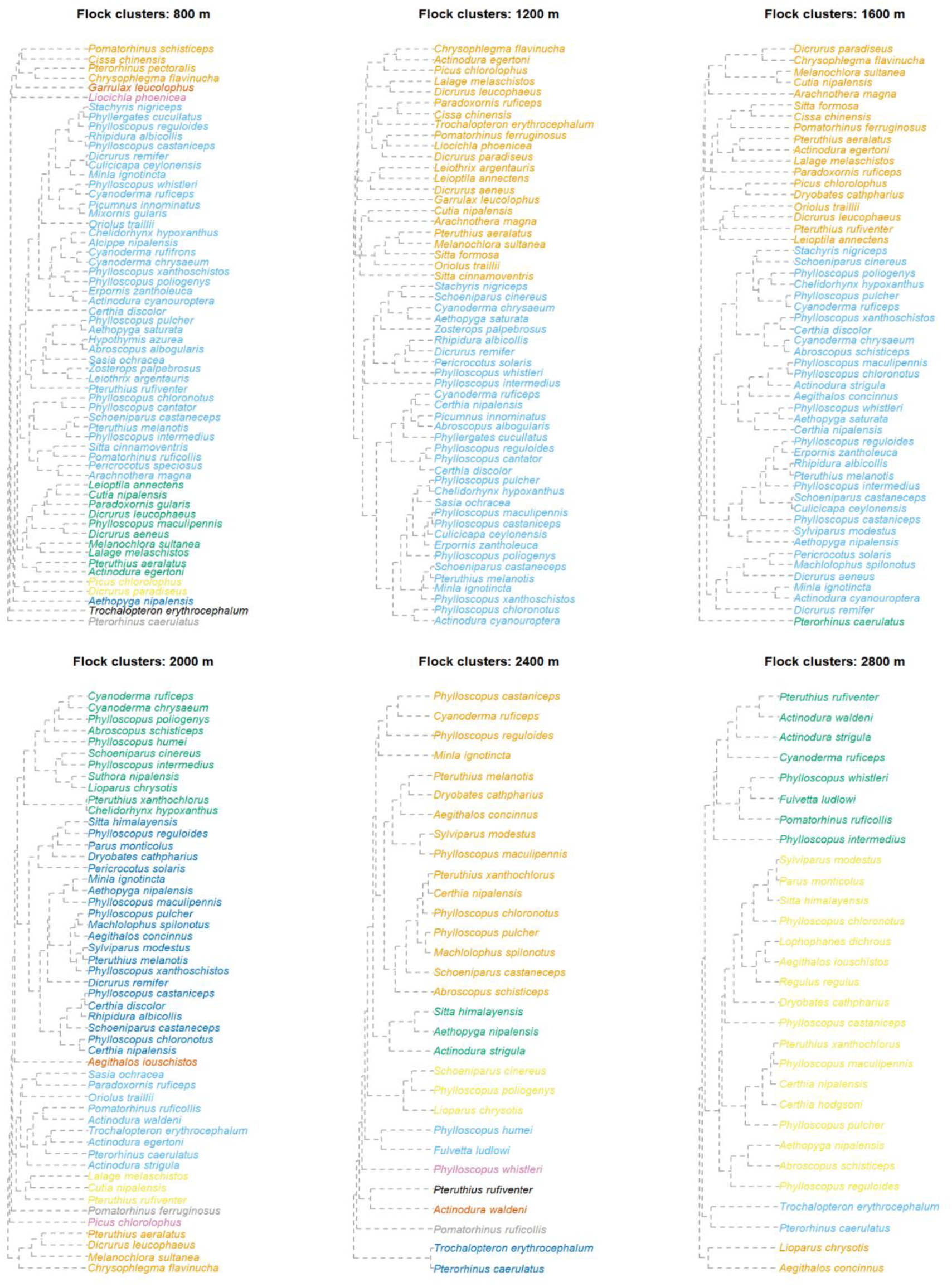
Cluster dendrograms of mixed-species bird flocks found at different elevations. Each tip represents a flocking species, and the colour of the tip represents the distinct MSF module to which each species belongs.

### Average elevational range overlap between flocking and non-flocking species

Mean elevational range overlap was found to be highest between (i) *within*-MSF module species pairs: 0.502 [95% CI = 0.493, 0.511], followed by (ii) *all* species pairs participating in MSFs: 0.390 [0.381, 0.399], (iii) species pairs in the overall species pool: 0.368 [0.365, 0.371] and the lowest for (iv) pairs of species not participating in MSFs: 0.360 [0.355,0.365] (Fig. 2).

**Figure 2.**
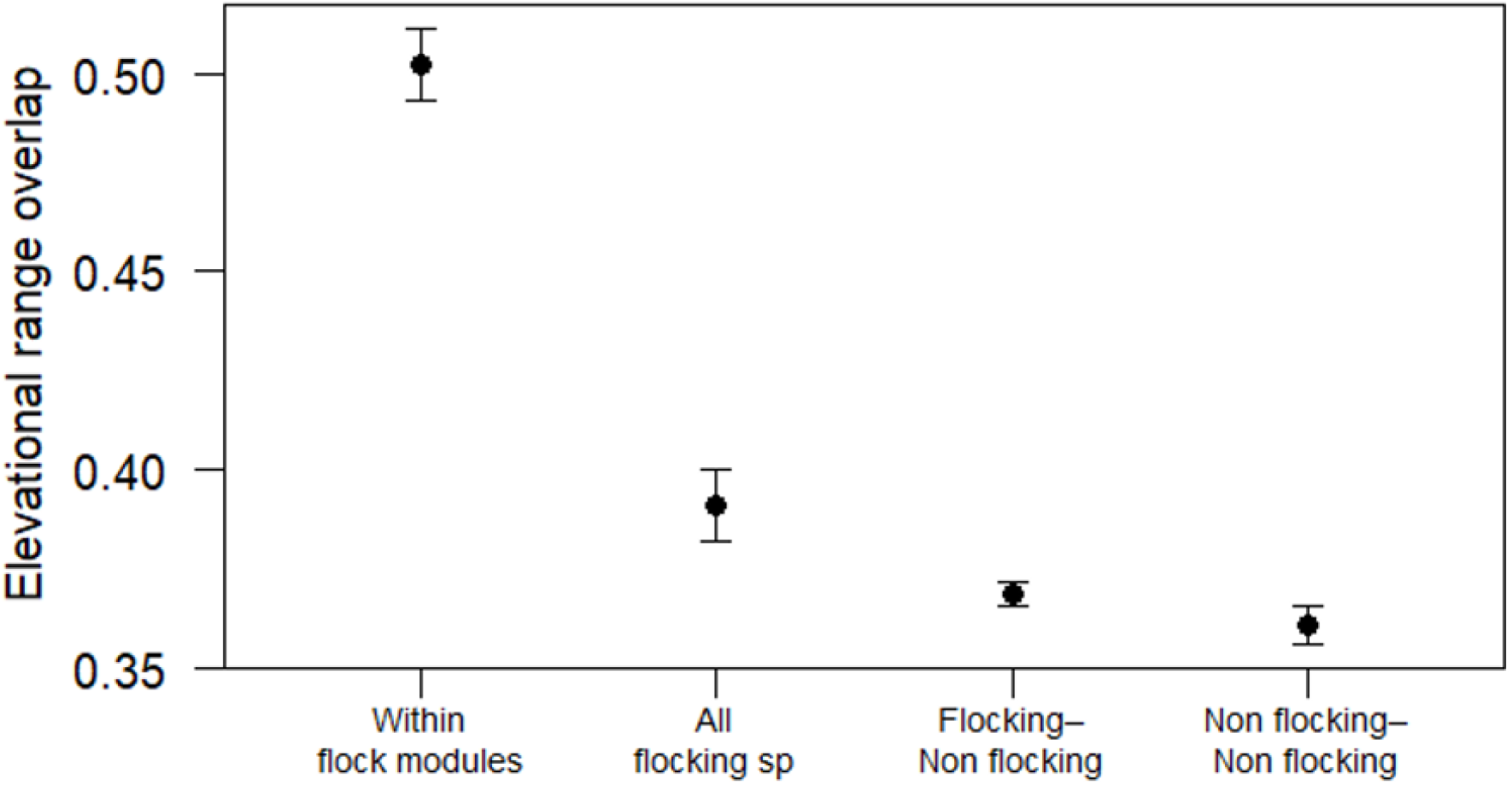
Mean elevational range overlap between pairs of: (i) flocking species belonging within unique MSF modules (ii) species which overall participate in MSFs, (iii) species both participating and not participating in MSFs, and (iv) species not participating in MSFs. Error bars represent 95% confidence intervals.

### Correlations between elevational range overlap and association strength

The relationship between association strength and degree of elevational range overlap for *all* species participating in MSFs was statistically significant and weakly positive (Pearson’s *r* = 0.30 [0.27, 0.33], *t*_3484_ = 18.69, p < 0.05; Fig. 3). Within-module correlations were generally higher (Pearson’s *r* ranging from 0.31 to 0.55), with the exception of two MSF modules: 1200m large-bodied and 1600m large-bodied MSFs with low *r* (=0.11 [-0.02, 0.24] and 0.09 [-0.07, 0.24]; Table S1; Fig 4).

**Figure 3.**
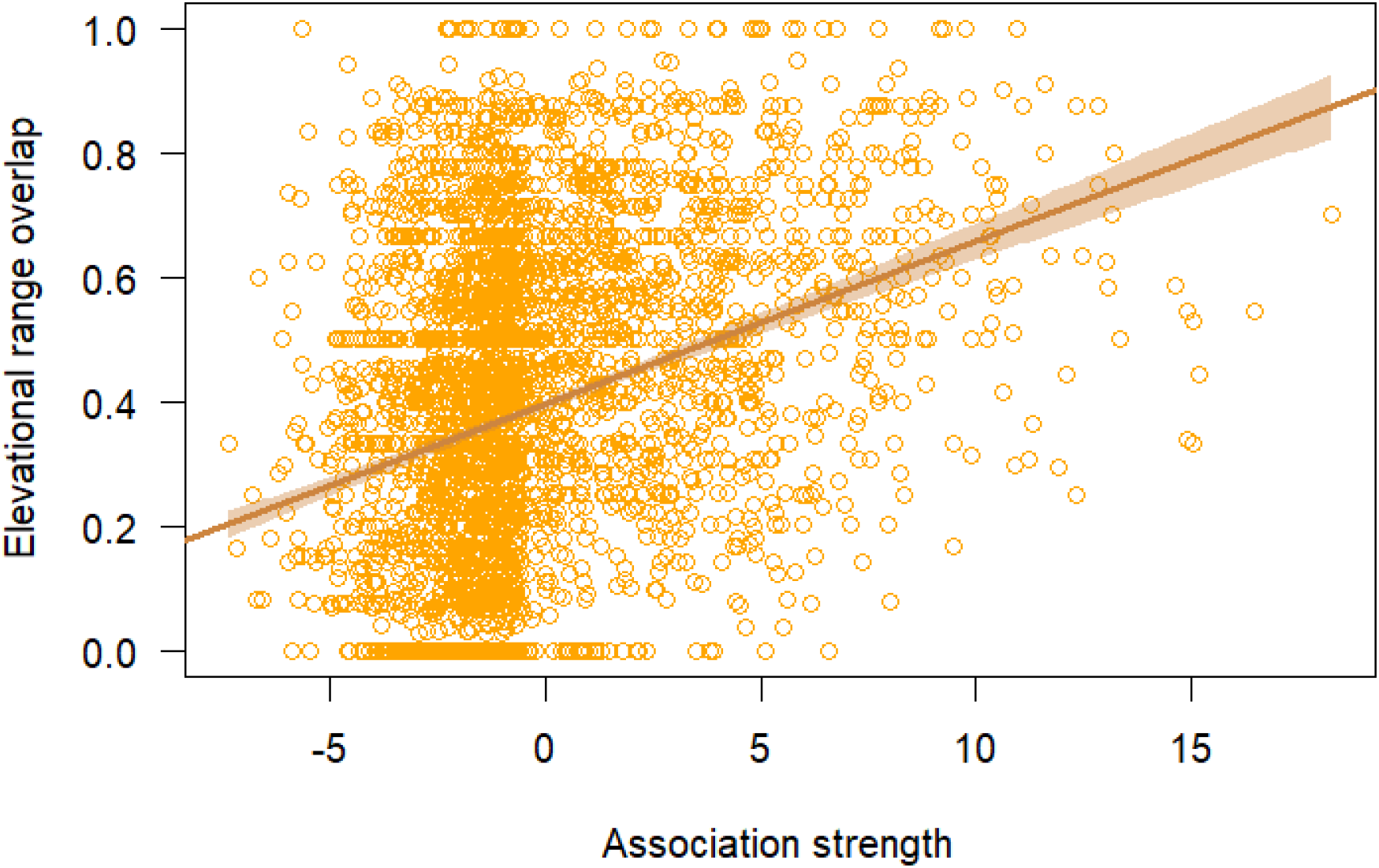
The relationship between association strength in MSFs and elevational range overlap values of pairs of all species that participate in MSFs.

**Figure 4.**
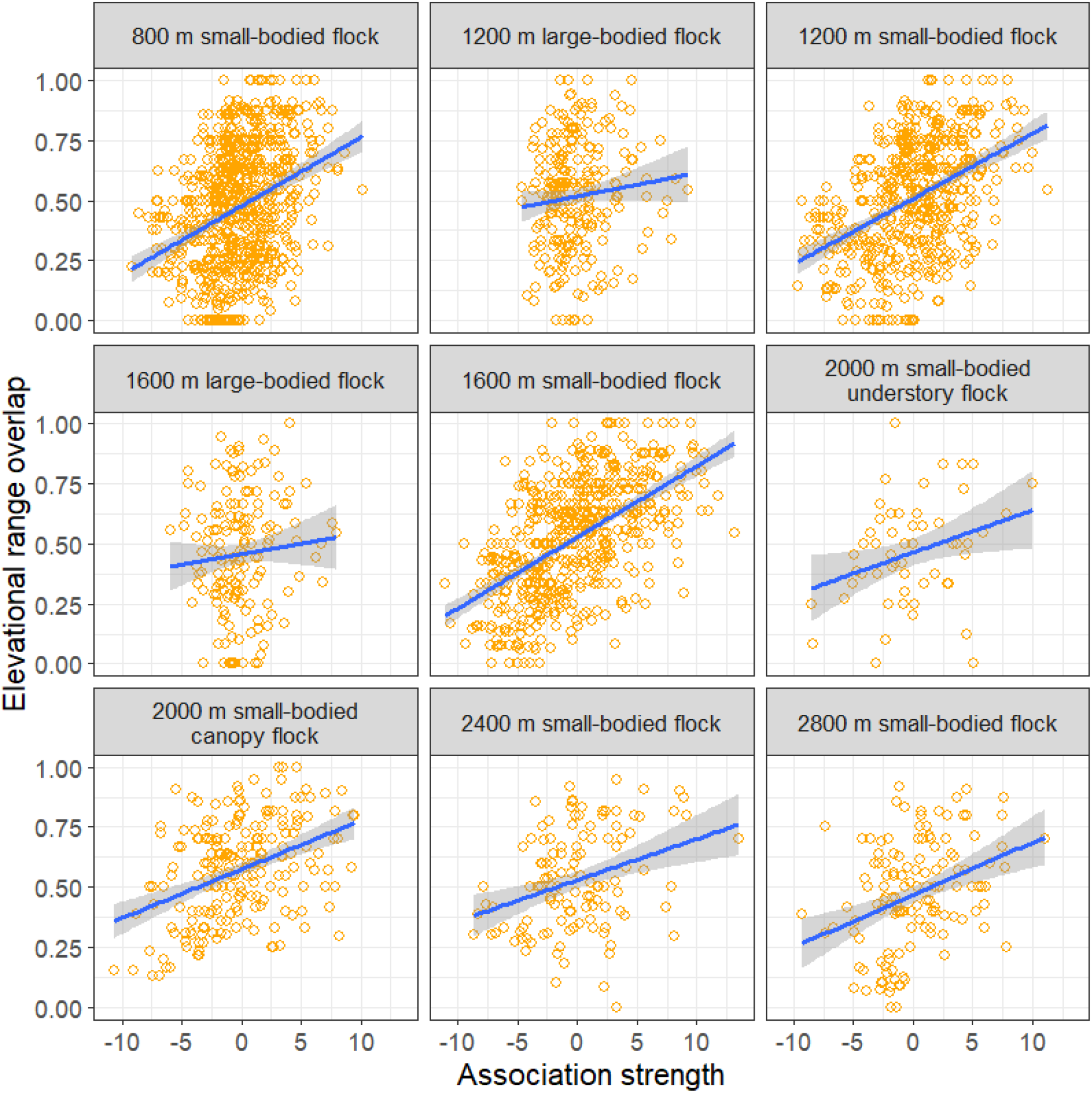
Correlations between association strength and elevational range overlap of species pairs within each unique MSF module. Each panel represents a unique MSF type; the title indicates elevation and the MSF type id. The locations are arranged in increasing order of elevation (800 to 2800m).

## Discussion

We show that on average, pairs of species that participating in mixed-species bird flocks (MSFs) showed a higher degree of winter elevational range overlap compared with those that do not, with range overlap values being higher for species pairs within compositionally distinct MSF modules (Fig. 2). Across all flocking species, pairwise association strength in MSFs was positively but weakly correlated with elevational range overlap. This relationship was consistently more strongly positive when examined separately for MSF participant pairs within compositionally distinct MSF modules (Figs. 3, 4, Table S1).

The higher degree of winter range overlap between species participating in MSFs is likely because of facilitative fitness benefits associated with participating in MSFs. During winter, when temperatures are lower and resources decline (including in the Himalayas and our study site; Ghosh et al. 2011, Menon et al. 2026), MSF participants can accrue fitness benefits such as increased foraging efficiency (through enhanced access to prey) and reduced predation risk (allowing for greater foraging time) through dilution and sentinel effects (Grubb Jr. 1987, Thiollay 1999, Mangini & Areta 2018). MSF participants can therefore co-occur in their realised niches, leading to high overlaps in spatial occurrence (Odling-Smee et al. 2003, Martínez et al. 2018). Unlike non-MSF participants, therefore, participating in MSFs with other species can promote range overlaps between participating species (Fig. 2) and consequently shape community structuring across elevational gradients. Using a different sampling protocol and a different dataset from the same elevational gradient sampled here, Mungee et al. (2025) hypothesise that winter bird communities are very likely structured by assembly processes other than environmental filtering and competition. While Mungee et al. (2025) speculate that this process could be elevational migration (that leads to the seasonal re-assortment of elevational ranges between summer and winter), facilitation within MSFs in winter is likely to be an important process determining winter elevational community assembly.

The positive (but statistically significant) relationship between pairwise association strength and range overlap for *all* flocking species (i.e., across-module; Fig. 3) irrespective of species-specific winter elevational ranges might be weak because even though species might co-occur elevationally, they do segregate into compositionally non-overlapping (or minimally overlapping) MSF modules (e.g., bamboo versus forest MSFs, understorey versus canopy MSFs; Srinivasan et al. 2012, Srinivasan et al. 2026). Therefore, two species belonging to different MSF modules but overlapping in elevational range, would have low association strengths but high range overlaps. For such species, similarities in elevational range might arise from similar requirements for macro-scale environmental characteristics such as temperature (i.e., comparable abiotic filtering; Pigot et al. 2010, Boucher-Lalonde et al. 2014, Elsen et al. 2017) followed by differences in—and adaptation to—more specific local scale habitat features (e.g., foraging on bamboo [Srinivasan et al. 2026] or preference for the forest canopy [Srinivasan et al. 2012, Montaño-Centellas & Jones 2021]). Indeed, identifying and separating MSF modules based on species composition and then testing for correlations between species pair-wise association strength (*within* these modules) and elevational range overlap consistently yields stronger relationships (Figs. 2, 4) than for all flocking species combined (Figs. 2, 3).

Within nine different MSF modules, correlations between association strength and elevational range overlaps were consistently stronger than for all species across modules. However, this correlation was weaker for the large-bodied MSFs at 1200 and 1600m (Fig. 4), in which species pairs tend to have high association strength but lower-than expected elevational range overlap. We suggest that species in these modules are likely involved in facultative facilitative interactions with different but functionally similar partner species in different parts of their elevational range. For instance, at our study site, *Cyanoderma chrysaeum* (Golden Babbler) flocks with different species at different elevations; *Alcippe nipalensis* (Nepal Fulvetta) at 800m ASL (n = 83 MSFs, association strength = 7.34, range overlap = 0.87), *Schoeniparus castaneceps* (Rufous-winged Fulvetta) between 800 and 2000m ASL (124, 7.02, 0.27), *Schoeniparus cinereus* (Yellow-throated Fulvetta) between 1200-2000m ASL (93, 4.07, 0.38) and *Lioparus chrysotis* (Golden-Breasted Fulvetta) at 2000m ASL in bamboo habitat (18, 1.07, 0.20). All these species are, however, intraspecifically gregarious and co-operatively breeding species. *C. chrysaeum* is therefore simply likely to join a gregarious species in MSFs regardless of species identity, thereby having a high association strength but low range overlap with each species (except with *A. nipalensis*). Intraspecific gregariousness is a well-known “attractive” trait of species that are preferentially joined by other species in MSFs (Sridhar & Shanker 2014), likely because such species provide antipredator benefits such as dilution and early warning (Bharadwaj et al. 2025).

Similar patterns are seen in the co-occurrence patterns of three low-elevation bamboo-specialist flocking species along latitude/longitude (i.e., in two-dimensional “classical” geographical range space). *Pomatorhinus ochraceiceps* (Red-billed Scimitar-Babbler) ranges from north-eastern India, through parts of Myanmar, Bangladesh, Laos, Vietnam and Thailand to southern China. In South Asia, *P. ochraceiceps* flocks with *Gampsorhynchus rufulus* (White-hooded Babbler) which occurs from south-eastern Nepal through the north-east Indian hill states, some regions in Bangladesh and China, to Myanmar. Where the distribution of *G. rufulus* ends, and the species is replaced by the functionally very similar congener *Gampsorhynchus torquatus* (Collared Babbler; distributed across Southeast Asia) *P. ochraceiceps* joins *G. torquatus* in MSFs (Billerman et al. 2026). Turnover in species participation in MSFs across large spatial gradients might nonetheless be associated with little turnover in the functional traits of MSF participants (i.e., conserved morphology, behaviour and/or phylogeny). A more detailed test of this hypothesis would include quantifying the joint abundance profiles of functionally similar species and overlaps with the abundance profiles of partner species along environmental gradients (Maitra et al. 2026).

Whereas our results are consistent with the hypothesis that associating strongly within facilitative MSFs can result in high range overlaps, the reverse might also be true – pairs of species that overlap in space because of similar abiotic and/or biotic requirements might also be more likely to associate within the same MSFs. Two scenarios can lead to the patterns we report. First, as we suggest, two species facilitate each other in MSFs such that mutual range expansion or constriction results in co-occurence in space (Stephan et al. 2021). This can occur when one species facilitates the persistence of the other species in areas (e.g., habitat, elevation) that would have been inhabitable in the absence of facilitative interactions (Martínez et al. 2018). For instance, alpine nurse plants alter adverse environmental conditions and enhance resource availability for other plant species (Callaway et al. 2002, Cavieres et al. 2014). Similarly, the geographical ranges of Neotropical obligate army ant-following birds closely coincide with those of army ants (Interiano et al. 2024). Second, abiotic filters might select for physiologically similar species, and such species then co-occur within similar geographic limits, followed by the subsequent coevolution of facilitative interactions between them (Luzuriaga et al. 2012).

While both scenarios are certainly possible, we suggest that the first is more likely based on the following evidence. Species belonging to distinct MSF types (Fig. 1) have much higher elevational range overlaps compared with that between all MSF species or all non-flocking species (Fig. 2). If all geographically overlapping species were indeed potentially free to participate in MSFs indiscriminately with any available partner species (second scenario in the paragraph above), within-module and across-module range overlaps should be comparable. Consequently, one would not expect multiple MSF modules at a particular elevation. However, it is now abundantly clear that sympatric MSF participants do indeed segregate strongly by habitat (Zhang et al. 2013, Mangini et al. 2023, Srinivasan et al. 2026), body size (Fig. 4) and vegetation stratum (King & Rappole 2001, Srinivasan et al. 2012, Montaño-Centellas & Jones 2021), and that species incur costs such as the need for activity matching in associations between functionally mismatched species (Hutto 1988, Sridhar & Guttal 2018). Because similar species are likely to benefit each other far better than functionally different species, they would be expected to co-occur in both MSFs (leading to high association strengths) and in geographical space (leading to high range overlaps). MSF modules are therefore very likely to reflect significant positive interactions between species within them, potentially influencing species’ realised niches (= ranges; Cai et al. 2020, Stephan et al. 2021).

Empirical work on MSFs (both experimental and observational) further supports the claim that facilitative interactions within MSFs can lead to changes in the realised niches and geographical ranges of participant species. For instance, the temporary removal of a highly effective sentinel species—which provides early warning against predators—from Neotropical MSFs causes other participant species to alter niches to forage in lower and denser forest strata (Martínez et al. 2018). In other words, many flocking species expand their niches to include the more open canopy in the presence of sentinel species (Martínez et al. 2018). In another study on MSF occurrence on islands of varying sizes in the Thousand Island Lake in China, smaller islands lack MSFs, likely because of the absence of intraspecifically gregarious species important for MSF cohesion (Martínez et al. 2024).

Species that naturally occur in large intraspecific groups and disproportionately lead MSFs require large islands, to persist; accordingly MSFs also occur only on large islands, showing how facilitative interactions are important in influencing occupancy, and by extension, range size (Martínez et al. 2024).

Conclusively establishing whether facilitation influences range overlaps or whether range overlaps promote the evolution of facilitation is likely to require experimentation. A possible set of such experiments can be:

i. quantifying overlap in thermal tolerance (i.e., upper and lower temperature limits within which an organism has normal physiological function) between pairs of species and examine correlations with their association strength,
ii. removing potential nuclear species (sensu Martínez et al. 2018) from their range edges to test how this might affect the fitness or distribution of associated flocking species (Dolby & Grubb 1998, Chan et al. 2019), and
iii. by examining MSF patterns with anthropogenic changes such as habitat fragmentation, which have inadvertently set up “experiments” such as the Thousand Island Lake system (Martínez et al. 2024).

Thus far, the vast majority of literature on the role of biotic interactions in setting species range limits has focused on competition, including classic papers such as Connell (1961), who showed how competition between closely related species of barnacles shapes the realised niches of species. At a broader scale, Diamond (1975) inferred that competition between ecologically similar bird species pairs might create a non-random, “checkerboard pattern” in the Bismarck Archipelago. More recent work demonstrates how interspecific interference competition limits elevational range overlap between closely related species (Jankowski et al. 2010, Freeman et al. 2019), with competition along hyperdiverse montane gradients leading to narrow elevational ranges with consequences for community structuring (Freeman et al. 2022). The role of facilitation in influencing community assembly, however, remains poorly studied.

Although experimental approaches have been employed to understand facilitation and its effect on species distributions (summarized in Stephan et al. 2021), these approaches largely examine specific species pairs, and extrapolating such approaches to entire communities, in which focal species pairs are embedded within complex interaction networks can be challenging. We attempt to address this gap by examining facilitation networks in MSFs and their correlation with geographical range overlaps. Taken together, facilitation between a diverse set of interacting species can structure communities across large gradients through spatial and temporal overlap between participants exchanging mutual benefits.

## Materials and Methods Study Area

We conducted this study in the Eastern Himalaya in Eaglenest Wildlife Sanctuary (EWS), West Kameng District, Arunachal Pradesh, India. The Eastern Himalaya is the second most biodiverse region on earth, after only the Amazon-western Andes elevational gradient (Grenyer et al. 2006). EWS is known for its rich biodiversity, especially for its avifauna. EWS has the second-highest richness of passerines and highest richness of insectivorous birds globally (Orme et al. 2005, Mungee et al. 2025). EWS spans an area of 218 sq km and has an altitudinal range of 500-3250m above sea level (ASL). Vegetation changes elevationally, from lowland evergreen and semi-evergreen forests (< 900m), broad-leaved subtropical forest dominated by oak species (800-1900m), broad-leaved temperate forest dominated by oaks, magnolias and rhododendrons (1800-2800m), coniferous temperate forest (2,800–3,200m) and bamboo stands found at all elevations (Choudhury 2003, Athreya 2006).

### MSF Sampling

We sampled mixed-species bird flocks (MSFs) in seven permanent plots across the elevational gradient in EWS at 800m, 1200m, 1600m, 2000m, 2400m and 2800m ASL during the winters (November-March) of 2019-20 and 2024-25. All plots were placed in old-growth forest. Sampling was done by walking transects in the plots from 0630 hrs to 1100 hrs and 1400 hrs to 1600 hrs, with the timings based on peak MSF activity in the region. Upon encountering an MSF, the following information was collected: (i) date, time and location of MSF, (ii) species composition of the MSF, (iii) number of individuals of each species, and (iv) forest stratum used by the MSF (understorey, midstory or canopy).

### Elevation ranges

Most species in the Eastern Himalaya migrate elevationally, occupying higher elevations in the summer (breeding season) and lower elevations in winter (Menon et al. 2025). For this study, we analysed the winter ranges of species occurring in EWS, because MSFs in the Eastern Himalaya only form in winter and do not persist in the breeding season. Therefore, participating in MSFs should influence winter species distribution and not the breeding elevational range. Winter range data for species that participate in MSFs / flocking species (used interchangeably) was collected from Srinivasan et al. (2018). The protocol followed in Srinivasan et al. (2018) was a modified line transect method from 1000m to 3000m ASL. Birds were sampled from 0500 hrs to 1030 hrs and then from 1700 hrs to 1900 hrs in October-December 2015 (Srinivasan et al. 2018). Each elevational transect was sampled three times. The species identities of insectivore and omnivore birds present in EWS but not participating in MSFs / non-flocking species (used interchangeably) were compiled from eBird (Sullivan et al. 2009) records from EWS curated using an application called MYNA (https://myna.stateofindiasbirds.in/). Winter range data for these species was also taken from Srinivasan et al. (2018) and further supplemented with data obtained from species accounts in Birds of the World (Billerman et al. 2026).

### Statistical Analysis Elevational range overlap

The elevational range distribution data consisted of the minimum and maximum elevation where each species is found in winter. Based on this information, we calculated the range overlap between all species pairs using the Jaccard’s similarity coefficient. Jaccard’s index calculates the intersection of the ranges of two species divided by the total range covered by both species:

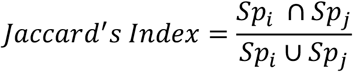

where *Sp*_*i*_ and *Sp*_*j*_ stands for the winter elevational range (lower elevational range limit to upper elevational range limit) of *i*^*th*^ and *j*^*th*^ species, respectively. For example, the Jaccard’s index value for *Sitta formosa* (Beautiful Nuthatch) and *Cutia nipalensis* (Himalayan Cutia) is 0.67, this value arising from *Sitta formosa* and *Cutia nipalensis* at our study site being found between 1200m-2000m and 1200m-2400m, respectively. In this case, the intersection, which is the common range in elevation where both are found, is (2000m − 1200m = 800m). And the union, which is the total range where at least one of the two species is found is (2400m − 1200m = 1200m). Putting the intersection and union values in the equation yields a Jaccard’s index of 800m/1200m = 0.67.

We then created a symmetric species × species matrix, in which cells contained Jaccard’s index values for each species pair. The index value can theoretically range from 0 (no range overlap between the two species) to 1 (complete range overlap between the two species).

### Average elevational range overlap between flocking and non-flocking species

To quantify range overlaps between different sets of species based on their participation in MSFs or not, we created three separate symmetric matrices: (i) range overlap information for only species participating in MSFs (see above), (ii) range overlaps for the entire pool of omnivorous and insectivorous birds (because MSFs are mostly formed by these two dietary guilds), and (iii) range overlaps for only those species which do not participate in MSFs. For each group, we calculated and visualised the average range overlap values along with 95% CI to visually assess differences in range overlap based on flocking behaviour.

### Network Analysis

To quantify association strengths between species pairs participating in MSFs, we first converted MSF participation data into a presence-absence matrix, in which each row represented a sampled individual MSF and each column was a unique species that participated in MSFs. This is a type of incidence matrix that stores information on whether a particular species was present (or not) in a particular MSF in binary format (presence = 1, absence = 0). Species which had an incidence score of ≤ 10 (i.e., occurred in ≤ 10 MSFs) and *Heterophasia pulchella* (Beautiful Sibia), *Dicaeum ignipectus* (Fire-breasted Flowerpecker), *Yuhina occipitalis* (Rufous-vented Yuhina), *Yuhina gularis* (Stripe-throated Yuhina), *Yuhina flavicollis* (Whiskered Yuhina) were not considered for further analysis, because these species might have joined MSFs accidentally and are not regular MSF participants (*sensu* Farley 2008). Based on the incidence matrix, we created an adjacency matrix by calculating the product of the incidence matrix with itself. This adjacency matrix contains information on the co-occurrences of each species pair, representing the number of MSFs in which the two species in each species pair co-occurred.

To control for species’ flocking propensities (Srinivasan et al. 2010) and to calculate association strength between species pairs, we used a null model approach. In this case, a null model is a randomized matrix of species co-occurrence built based on random expectation arising from observed species-specific frequency of participation in MSFs. Our preferred choice of null model was the one which held row and column totals identical to that observed in the actual data (Connor & Simberloff 1979). Row totals represent the number of species observed in each MSF and column totals represent the observed number of MSFs that each species participated in. We used the curveball function in package {Incidentally} (Neal 2022) in the statistical and programming software R (R Core Team 2025) to construct the null matrices. Based on randomized reorganization of the original co-occurrence matrix, we constructed 1,000 null matrices and stored these in an array.

To quantify association strength between species’ pairs, we calculated an index (Srinivasan et al. 2010) based on the formula:

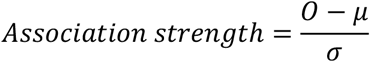

where *O* is the observed number of co-occurrence between two species, *μ* and *σ* are, respectively, the average co-occurrence value and standard deviation calculated across the 1,000 null matrices, respectively. A positive score implies greater association between two species than expected at random, and a negative score implies lower association than expected by random between the two species, whereas a score of zero would imply that there is no significant interaction (or a neutral interaction) between the two species (Sridhar et al. 2012).

### Correlation between elevational range overlap and association strength

#### Pearson’s product-moment correlation

To further quantify correlations between association score and elevational range overlap (with each data point being a species pair) we ran a Pearson’s product-moment correlation using the cor.test() function in R program (R Core Team 2025).

#### Characterisation of unique MSF modules and checking elevational range overlap within them

We predicted that average range overlap will be higher and there will be a stronger correlation between association strength and range overlap between species pairs belonging to a unique MSF type. During sampling, we observed that there were distinct MSF modules with differing species composition, with little overlap in species participation across modules. This has been repeatedly noted from previous studies from the same region (Srinivasan et al. 2010, 2012, Bharadwaj et al. 2025, Srinivasan et al. 2026, Gupta et al. 2026).

To account for (a) the fact that a large number of species participate in MSFs, but (b) in well-demarcated MSF “modules” with little overlap in species composition, and therefore (c) species across modules are highly unlikely to facilitate each other, we used the cluster_walktrap() function in package {igraph} in R program (R Core Team 2025) to separate different MSF modules based on species composition. Through short random walks, this function identifies densely connected sets of species (nodes) in “subgraphs”, such that the random walker tends to stay “trapped” within modules with highly connected nodes, and sparsely connected to nodes of other subgraphs (Malliaros & Vazirgiannis 2013, Blašković et al. 2025). In our analysis, each subgraph consisted of a distinct set of flocking species and were classified as a unique “MSF type”. From each sampled elevation, we extracted only those MSF types which had more than 10 unique species within them.

Unlike the “global” analysis of MSF participating species outlined above, we further calculated elevational range overlap between species pairs separately within each MSF module using Jaccard’s index. After which we calculated and visualized the average overlap values and the 95% CI among all the MSF modules.

#### Correlation between association strength and range overlap within MSF modules

We calculated association strength between species pairs separately within each MSF module (using the same analytical approach mentioned in the network analysis section above) while the null model was constructed using an alternative framework. Again, we considered four possible null modelling approaches. These were:

i. A null model in which species, while joining others, were not bound by any elevational range limits and are free to associate with any flocking species found along the entire elevational gradient.
ii. A null model in which species, while joining MSFs, are bound by the elevational ranges within which they are generally found but are free to associate with any flocking species also found at the in some part of its elevational range.
iii. A null model in which species, while joining MSFs, are not bound by any elevational range limits but are restricted to associate with only those species which are members of that unique MSF type.
iv. A null model in which species, while joining MSFs, are both bound by elevational range limits and choice of partner species based on which MSF type in which it is found.

We selected the third null model because it seemed the most appropriate based on biological considerations. Because we had *a priori* information on which species flocked with which others (e.g., Srinivasan et al. 2010, 2012, Bharadwaj et al. 2025, Srinivasan et al. 2026, Gupta et al. 2026), using only this subset for randomisation is likely more accurate than using the entire flocking species pool. Moreover, because we aimed to understand the degree of correlation between association strength and elevational range overlap, constraining the overall flocking species pool by elevational range would likely defeat the purpose of detecting association strength-range overlap relationships.

Finally, to find the correlation between association strength and range overlap between species pairs within an MSF module, we employed Pearson’s product-moment correlation and visualised the pattern using package {ggplot2} in R program (R Core Team 2025).

## Acknowledgments

We thank the Ministry of Environment, Forest and Climate Change, Government of India, Anusandhan National Research Foundation, Government of India, the Department of Biotechnology, Government of India and the Infosys Foundation for funding support. We are grateful to the Arunachal Pradesh Forest Department and the Shergaon Forest Division for their support of this project and for research permits to conduct studies in the Eaglenest Wildlife Sanctuary, Arunachal Pradesh. We thank The Management Committee of the Singchung Bugun Village Community Reserve, especially Mr Khandu Glow for their support, hospitality and permission to work within the Bugun community lands of Singchung village.

