## Supplementary Table 1 and 2 for "Positive relationships between interspecific association strength and elevational range overlap highlight the role of facilitation in community assembly"

Sarthak Mondal

Tables

Table S1. Pearson’s correlation results between association strength and elevational range overlap values of pairs of flocking species within each unique MSF type. Note that there might be multiple distinct modules at some elevations (e.g., 1200m ASL).

| **Sl. No** | **Elevation** | **MSF module** | **Pearson’s correlation** | **t-statistics** | **p-value** |
| --- | --- | --- | --- | --- | --- |
| 1. | 800m | small-bodied | 0.33 [0.26, 0.39] | t_(739)_ = 9.42 | <0.05 |
| 2. | 1200m | large-bodied | 0.11 [-0.02, 0.24] | t_(229)_ = 1.70 | 0.09 |
| 3. | 1200m | small-bodied | 0.43 [0.36, 0.50] | t_(494)_ = 10.70 | <0.05 |
| 4. | 1600m | large-bodied | 0.09 [-0.07, 0.24] | t_(151)_ = 1.10 | 0.27 |
| 5. | 1600m | small-bodied | 0.55 [0.49, 0.61] | t_(526)_ = 15.20 | <0.05 |
| 6. | 2000m | small-bodied understory | 0.31 [0.04, 0.53] | t_(53)_ = 2.34 | 0.02 |
| 7. | 2000m | small-bodied canopy | 0.40 [0.28, 0.51] | t_(208)_ = 6.33 | <0.05 |
| 8. | 2400m | small-bodied | 0.32 [0.15, 0.47] | t_(118)_ = 3.69 | <0.05 |
| 9. | 2800m | small-bodied | 0.34 [0.19, 0.48] | t_(134)_ = 4.22 | <0.05 |

Table S2. Summary of the different MSF modules having more than 10 constituent species.

| **Sl. No.** | **Elevation** | **MSF module** | **Species (Scientific Name)** | **Species (Common Name)** |
| --- | --- | --- | --- | --- |
| 1 | 800m | small-bodied | *Abroscopus albogularis* | Rufous-faced Warbler |
| 2 | 800m | small-bodied | *Actinodura cyanouroptera* | Blue-winged Minla |
| 3 | 800m | small-bodied | *Aethopyga saturata* | Black-throated Sunbird |
| 4 | 800m | small-bodied | *Alcippe nipalensis* | Nepal Fulvetta |
| 5 | 800m | small-bodied | *Arachnothera magna* | Streaked Spiderhunter |
| 6 | 800m | small-bodied | *Certhia discolor* | Sikkim Treecreeper |
| 7 | 800m | small-bodied | *Chelidorhynx hypoxanthus* | Yellow-bellied Fairy-Fantail |
| 8 | 800m | small-bodied | *Culicicapa ceylonensis* | Gray-headed Canary-Flycatcher |
| 9 | 800m | small-bodied | *Cyanoderma chrysaeum* | Golden Babbler |
| 10 | 800m | small-bodied | *Cyanoderma ruficeps* | Rufous-capped Babbler |
| 11 | 800m | small-bodied | *Cyanoderma rufifrons* | Rufous-fronted Babbler |
| 12 | 800m | small-bodied | *Dicrurus remifer* | Lesser Racket-tailed Drongo |
| 13 | 800m | small-bodied | *Erpornis zantholeuca* | White-bellied Erpornis |
| 14 | 800m | small-bodied | *Hypothymis azurea* | Black-naped Monarch |
| 15 | 800m | small-bodied | *Leiothrix argentauris* | Silver-eared Mesia |
| 16 | 800m | small-bodied | *Minla ignotincta* | Red-tailed Minla |
| 17 | 800m | small-bodied | *Mixornis gularis* | Pin-striped Tit-Babbler |
| 18 | 800m | small-bodied | *Oriolus traillii* | Maroon Oriole |
| 19 | 800m | small-bodied | *Pericrocotus speciosus* | Scarlet Minivet |
| 20 | 800m | small-bodied | *Phyllergates cucullatus* | Mountain Tailorbird |
| 21 | 800m | small-bodied | *Phylloscopus cantator* | Yellow-vented Warbler |
| 22 | 800m | small-bodied | *Phylloscopus castaniceps* | Chestnut-crowned Warbler |
| 23 | 800m | small-bodied | *Phylloscopus chloronotus* | Lemon-rumped Warbler |
| 24 | 800m | small-bodied | *Phylloscopus intermedius* | White-spectacled Warbler |
| 25 | 800m | small-bodied | *Phylloscopus poliogenys* | Gray-cheeked Warbler |
| 26 | 800m | small-bodied | *Phylloscopus pulcher* | Buff-barred Warbler |
| 27 | 800m | small-bodied | *Phylloscopus reguloides* | Blyth's Leaf Warbler |
| 28 | 800m | small-bodied | *Phylloscopus whistleri* | Whistler's Warbler |
| 29 | 800m | small-bodied | *Phylloscopus xanthoschistos* | Gray-hooded Warbler |
| 30 | 800m | small-bodied | *Picumnus innominatus* | Speckled Piculet |
| 31 | 800m | small-bodied | *Pomatorhinus ruficollis* | Streak-breasted Scimitar-Babbler |
| 32 | 800m | small-bodied | *Pteruthius melanotis* | Black-eared Shrike-Babbler |
| 33 | 800m | small-bodied | *Pteruthius rufiventer* | Black-headed Shrike-Babbler |
| 34 | 800m | small-bodied | *Rhipidura albicollis* | White-throated Fantail |
| 35 | 800m | small-bodied | *Sasia ochracea* | White-browed Piculet |
| 36 | 800m | small-bodied | *Schoeniparus castaneceps* | Rufous-winged Fulvetta |
| 37 | 800m | small-bodied | *Sitta cinnamoventris* | Chestnut-bellied Nuthatch |
| 38 | 800m | small-bodied | *Stachyris nigriceps* | Gray-throated Babbler |
| 39 | 800m | small-bodied | *Zosterops palpebrosus* | Indian White-eye |
| 1 | 1200m | large-bodied | *Actinodura egertoni* | Rusty-fronted Barwing |
| 2 | 1200m | large-bodied | *Arachnothera magna* | Streaked Spiderhunter |
| 3 | 1200m | large-bodied | *Chrysophlegma flavinucha* | Greater Yellownape |
| 4 | 1200m | large-bodied | *Cissa chinensis* | Common Green-Magpie |
| 5 | 1200m | large-bodied | *Cutia nipalensis* | Himalayan Cutia |
| 6 | 1200m | large-bodied | *Dicrurus aeneus* | Bronzed Drongo |
| 7 | 1200m | large-bodied | *Dicrurus leucophaeus* | Ashy Drongo |
| 8 | 1200m | large-bodied | *Dicrurus paradiseus* | Greater Racket-tailed Drongo |
| 9 | 1200m | large-bodied | *Garrulax leucolophus* | White-crested Laughingthrush |
| 10 | 1200m | large-bodied | *Lalage melaschistos* | Black-winged Cuckooshrike |
| 11 | 1200m | large-bodied | *Leioptila annectens* | Rufous-backed Sibia |
| 12 | 1200m | large-bodied | *Leiothrix argentauris* | Silver-eared Mesia |
| 13 | 1200m | large-bodied | *Liocichla phoenicea* | Red-faced Liocichla |
| 14 | 1200m | large-bodied | *Melanochlora sultanea* | Sultan Tit |
| 15 | 1200m | large-bodied | *Oriolus traillii* | Maroon Oriole |
| 16 | 1200m | large-bodied | *Paradoxornis ruficeps* | White-breasted Parrotbill |
| 17 | 1200m | large-bodied | *Picus chlorolophus* | Lesser Yellownape |
| 18 | 1200m | large-bodied | *Pomatorhinus ferruginosus* | Black-crowned Scimitar-Babbler |
| 19 | 1200m | large-bodied | *Pteruthius aeralatus* | White-browed Shrike-Babbler |
| 20 | 1200m | large-bodied | *Sitta cinnamoventris* | Chestnut-bellied Nuthatch |
| 21 | 1200m | large-bodied | *Sitta formosa* | Beautiful Nuthatch |
| 22 | 1200m | large-bodied | *Trochalopteron erythrocephalum* | Chestnut-crowned Laughingthrush |
| 1 | 1200m | small-bodied | *Abroscopus albogularis* | Rufous-faced Warbler |
| 2 | 1200m | small-bodied | *Actinodura cyanouroptera* | Blue-winged Minla |
| 3 | 1200m | small-bodied | *Aethopyga saturata* | Black-throated Sunbird |
| 4 | 1200m | small-bodied | *Certhia discolor* | Sikkim Treecreeper |
| 5 | 1200m | small-bodied | *Certhia nipalensis* | Rusty-flanked Treecreeper |
| 6 | 1200m | small-bodied | *Chelidorhynx hypoxanthus* | Yellow-bellied Fairy-Fantail |
| 7 | 1200m | small-bodied | *Culicicapa ceylonensis* | Gray-headed Canary-Flycatcher |
| 8 | 1200m | small-bodied | *Cyanoderma chrysaeum* | Golden Babbler |
| 9 | 1200m | small-bodied | *Cyanoderma ruficeps* | Rufous-capped Babbler |
| 10 | 1200m | small-bodied | *Dicrurus remifer* | Lesser Racket-tailed Drongo |
| 11 | 1200m | small-bodied | *Erpornis zantholeuca* | White-bellied Erpornis |
| 12 | 1200m | small-bodied | *Minla ignotincta* | Red-tailed Minla |
| 13 | 1200m | small-bodied | *Pericrocotus solaris* | Gray-chinned Minivet |
| 14 | 1200m | small-bodied | *Phyllergates cucullatus* | Mountain Tailorbird |
| 15 | 1200m | small-bodied | *Phylloscopus cantator* | Yellow-vented Warbler |
| 16 | 1200m | small-bodied | *Phylloscopus castaniceps* | Chestnut-crowned Warbler |
| 17 | 1200m | small-bodied | *Phylloscopus chloronotus* | Lemon-rumped Warbler |
| 18 | 1200m | small-bodied | *Phylloscopus intermedius* | White-spectacled Warbler |
| 19 | 1200m | small-bodied | *Phylloscopus maculipennis* | Ashy-throated Warbler |
| 20 | 1200m | small-bodied | *Phylloscopus poliogenys* | Gray-cheeked Warbler |
| 21 | 1200m | small-bodied | *Phylloscopus pulcher* | Buff-barred Warbler |
| 22 | 1200m | small-bodied | *Phylloscopus reguloides* | Blyth's Leaf Warbler |
| 23 | 1200m | small-bodied | *Phylloscopus whistleri* | Whistler's Warbler |
| 24 | 1200m | small-bodied | *Phylloscopus xanthoschistos* | Gray-hooded Warbler |
| 25 | 1200m | small-bodied | *Picumnus innominatus* | Speckled Piculet |
| 26 | 1200m | small-bodied | *Pteruthius melanotis* | Black-eared Shrike-Babbler |
| 27 | 1200m | small-bodied | *Rhipidura albicollis* | White-throated Fantail |
| 28 | 1200m | small-bodied | *Sasia ochracea* | White-browed Piculet |
| 29 | 1200m | small-bodied | *Schoeniparus castaneceps* | Rufous-winged Fulvetta |
| 30 | 1200m | small-bodied | *Schoeniparus cinereus* | Yellow-throated Fulvetta |
| 31 | 1200m | small-bodied | *Stachyris nigriceps* | Gray-throated Babbler |
| 32 | 1200m | small-bodied | *Zosterops palpebrosus* | Indian White-eye |
| 1 | 1600m | large-bodied | *Actinodura egertoni* | Rusty-fronted Barwing |
| 2 | 1600m | large-bodied | *Arachnothera magna* | Streaked Spiderhunter |
| 3 | 1600m | large-bodied | *Chrysophlegma flavinucha* | Greater Yellownape |
| 4 | 1600m | large-bodied | *Cissa chinensis* | Common Green-Magpie |
| 5 | 1600m | large-bodied | *Cutia nipalensis* | Himalayan Cutia |
| 6 | 1600m | large-bodied | *Dicrurus leucophaeus* | Ashy Drongo |
| 7 | 1600m | large-bodied | *Dicrurus paradiseus* | Greater Racket-tailed Drongo |
| 8 | 1600m | large-bodied | *Dryobates cathpharius* | Crimson-naped Woodpecker |
| 9 | 1600m | large-bodied | *Lalage melaschistos* | Black-winged Cuckooshrike |
| 10 | 1600m | large-bodied | *Leioptila annectens* | Rufous-backed Sibia |
| 11 | 1600m | large-bodied | *Melanochlora sultanea* | Sultan Tit |
| 12 | 1600m | large-bodied | *Oriolus traillii* | Maroon Oriole |
| 13 | 1600m | large-bodied | *Paradoxornis ruficeps* | White-breasted Parrotbill |
| 14 | 1600m | large-bodied | *Picus chlorolophus* | Lesser Yellownape |
| 15 | 1600m | large-bodied | *Pomatorhinus ferruginosus* | Black-crowned Scimitar-Babbler |
| 16 | 1600m | large-bodied | *Pteruthius aeralatus* | White-browed Shrike-Babbler |
| 17 | 1600m | large-bodied | *Pteruthius rufiventer* | Black-headed Shrike-Babbler |
| 18 | 1600m | large-bodied | *Sitta formosa* | Beautiful Nuthatch |
| 1 | 1600m | small-bodied | *Abroscopus schisticeps* | Black-faced Warbler |
| 2 | 1600m | small-bodied | *Actinodura cyanouroptera* | Blue-winged Minla |
| 3 | 1600m | small-bodied | *Actinodura strigula* | Chestnut-tailed Minla |
| 4 | 1600m | small-bodied | *Aegithalos concinnus* | Black-throated Tit |
| 5 | 1600m | small-bodied | *Aethopyga nipalensis* | Green-tailed Sunbird |
| 6 | 1600m | small-bodied | *Aethopyga saturata* | Black-throated Sunbird |
| 7 | 1600m | small-bodied | *Certhia discolor* | Sikkim Treecreeper |
| 8 | 1600m | small-bodied | *Certhia nipalensis* | Rusty-flanked Treecreeper |
| 9 | 1600m | small-bodied | *Chelidorhynx hypoxanthus* | Yellow-bellied Fairy-Fantail |
| 10 | 1600m | small-bodied | *Culicicapa ceylonensis* | Gray-headed Canary-Flycatcher |
| 11 | 1600m | small-bodied | *Cyanoderma chrysaeum* | Golden Babbler |
| 12 | 1600m | small-bodied | *Cyanoderma ruficeps* | Rufous-capped Babbler |
| 13 | 1600m | small-bodied | *Dicrurus aeneus* | Bronzed Drongo |
| 14 | 1600m | small-bodied | *Dicrurus remifer* | Lesser Racket-tailed Drongo |
| 15 | 1600m | small-bodied | *Erpornis zantholeuca* | White-bellied Erpornis |
| 16 | 1600m | small-bodied | *Machlolophus spilonotus* | Yellow-cheeked Tit |
| 17 | 1600m | small-bodied | *Minla ignotincta* | Red-tailed Minla |
| 18 | 1600m | small-bodied | *Pericrocotus solaris* | Gray-chinned Minivet |
| 19 | 1600m | small-bodied | *Phylloscopus castaniceps* | Chestnut-crowned Warbler |
| 20 | 1600m | small-bodied | *Phylloscopus chloronotus* | Lemon-rumped Warbler |
| 21 | 1600m | small-bodied | *Phylloscopus intermedius* | White-spectacled Warbler |
| 22 | 1600m | small-bodied | *Phylloscopus maculipennis* | Ashy-throated Warbler |
| 23 | 1600m | small-bodied | *Phylloscopus poliogenys* | Gray-cheeked Warbler |
| 24 | 1600m | small-bodied | *Phylloscopus pulcher* | Buff-barred Warbler |
| 25 | 1600m | small-bodied | *Phylloscopus reguloides* | Blyth's Leaf Warbler |
| 26 | 1600m | small-bodied | *Phylloscopus whistleri* | Whistler's Warbler |
| 27 | 1600m | small-bodied | *Phylloscopus xanthoschistos* | Gray-hooded Warbler |
| 28 | 1600m | small-bodied | *Pteruthius melanotis* | Black-eared Shrike-Babbler |
| 29 | 1600m | small-bodied | *Rhipidura albicollis* | White-throated Fantail |
| 30 | 1600m | small-bodied | *Schoeniparus castaneceps* | Rufous-winged Fulvetta |
| 31 | 1600m | small-bodied | *Schoeniparus cinereus* | Yellow-throated Fulvetta |
| 32 | 1600m | small-bodied | *Stachyris nigriceps* | Gray-throated Babbler |
| 33 | 1600m | small-bodied | *Sylviparus modestus* | Yellow-browed Tit |
| 1 | 2000m | small-bodied understory | *Abroscopus schisticeps* | Black-faced Warbler |
| 2 | 2000m | small-bodied understory | *Aegithalos concinnus* | Black-throated Parrotbill |
| 3 | 2000m | small-bodied understory | *Chelidorhynx hypoxanthus* | Yellow-bellied Fairy-Fantail |
| 4 | 2000m | small-bodied understory | *Cyanoderma chrysaeum* | Golden Babbler |
| 5 | 2000m | small-bodied understory | *Cyanoderma ruficeps* | Rufous-capped Babbler |
| 6 | 2000m | small-bodied understory | *Lioparus chrysotis* | Golden-breasted Fulvetta |
| 7 | 2000m | small-bodied understory | *Phylloscopus humei* | Hume's Warbler |
| 8 | 2000m | small-bodied understory | *Phylloscopus intermedius* | White-spectacled Warbler |
| 9 | 2000m | small-bodied understory | *Phylloscopus poliogenys* | Gray-cheeked Warbler |
| 10 | 2000m | small-bodied understory | *Pteruthius xanthochlorus* | Green Shrike-Babbler |
| 11 | 2000m | small-bodied understory | *Schoeniparus cinereus* | Yellow-throated Fulvetta |
| 1 | 2000m | small-bodied canopy | *Aegithalos concinnus* | Black-throated Tit |
| 2 | 2000m | small-bodied canopy | *Aethopyga nipalensis* | Green-tailed Sunbird |
| 3 | 2000m | small-bodied canopy | *Certhia discolor* | Sikkim Treecreeper |
| 4 | 2000m | small-bodied canopy | *Certhia nipalensis* | Rusty-flanked Treecreeper |
| 5 | 2000m | small-bodied canopy | *Dicrurus remifer* | Lesser Racket-tailed Drongo |
| 6 | 2000m | small-bodied canopy | *Dryobates cathpharius* | Crimson-naped Woodpecker |
| 7 | 2000m | small-bodied canopy | *Machlolophus spilonotus* | Yellow-cheeked Tit |
| 8 | 2000m | small-bodied canopy | *Minla ignotincta* | Red-tailed Minla |
| 9 | 2000m | small-bodied canopy | *Parus monticolus* | Green-backed Tit |
| 10 | 2000m | small-bodied canopy | *Pericrocotus solaris* | Gray-chinned Minivet |
| 11 | 2000m | small-bodied canopy | *Phylloscopus castaniceps* | Chestnut-crowned Warbler |
| 12 | 2000m | small-bodied canopy | *Phylloscopus chloronotus* | Lemon-rumped Warbler |
| 13 | 2000m | small-bodied canopy | *Phylloscopus maculipennis* | Ashy-throated Warbler |
| 14 | 2000m | small-bodied canopy | *Phylloscopus pulcher* | Buff-barred Warbler |
| 15 | 2000m | small-bodied canopy | *Phylloscopus reguloides* | Blyth's Leaf Warbler |
| 16 | 2000m | small-bodied canopy | *Phylloscopus xanthoschistos* | Gray-hooded Warbler |
| 17 | 2000m | small-bodied canopy | *Pteruthius melanotis* | Black-eared Shrike-Babbler |
| 18 | 2000m | small-bodied canopy | *Rhipidura albicollis* | White-throated Fantail |
| 19 | 2000m | small-bodied canopy | *Schoeniparus castaneceps* | Rufous-winged Fulvetta |
| 20 | 2000m | small-bodied canopy | *Sitta himalayensis* | White-tailed Nuthatch |
| 21 | 2000m | small-bodied canopy | *Sylviparus modestus* | Yellow-browed Tit |
| 1 | 2400m | small-bodied | *Abroscopus schisticeps* | Black-faced Warbler |
| 2 | 2400m | small-bodied | *Aegithalos concinnus* | Black-throated Tit |
| 3 | 2400m | small-bodied | *Certhia nipalensis* | Rusty-flanked Treecreeper |
| 4 | 2400m | small-bodied | *Cyanoderma ruficeps* | Rufous-capped Babbler |
| 5 | 2400m | small-bodied | *Dryobates cathpharius* | Crimson-naped Woodpecker |
| 6 | 2400m | small-bodied | *Machlolophus spilonotus* | Yellow-cheeked Tit |
| 7 | 2400m | small-bodied | *Minla ignotincta* | Red-tailed Minla |
| 8 | 2400m | small-bodied | *Phylloscopus castaniceps* | Chestnut-crowned Warbler |
| 9 | 2400m | small-bodied | *Phylloscopus chloronotus* | Lemon-rumped Warbler |
| 10 | 2400m | small-bodied | *Phylloscopus maculipennis* | Ashy-throated Warbler |
| 11 | 2400m | small-bodied | *Phylloscopus pulcher* | Buff-barred Warbler |
| 12 | 2400m | small-bodied | *Phylloscopus reguloides* | Blyth's Leaf Warbler |
| 13 | 2400m | small-bodied | *Pteruthius melanotis* | Black-eared Shrike-Babbler |
| 14 | 2400m | small-bodied | *Pteruthius xanthochlorus* | Green Shrike-Babbler |
| 15 | 2400m | small-bodied | *Schoeniparus castaneceps* | Rufous-winged Fulvetta |
| 16 | 2400m | small-bodied | *Sylviparus modestus* | Yellow-browed Tit |
| 1 | 2800m | small-bodied | *Abroscopus schisticeps* | Black-faced Warbler |
| 2 | 2800m | small-bodied | *Aegithalos iouschistos* | Black-browed Tit |
| 3 | 2800m | small-bodied | *Aethopyga nipalensis* | Green-tailed Sunbird |
| 4 | 2800m | small-bodied | *Certhia hodgsoni* | Hodgson's Treecreeper |
| 5 | 2800m | small-bodied | *Certhia nipalensis* | Rusty-flanked Treecreeper |
| 6 | 2800m | small-bodied | *Dryobates cathpharius* | Crimson-naped Woodpecker |
| 7 | 2800m | small-bodied | *Lophophanes dichrous* | Gray-crested Tit |
| 8 | 2800m | small-bodied | *Parus monticolus* | Green-backed Tit |
| 9 | 2800m | small-bodied | *Phylloscopus castaniceps* | Chestnut-crowned Warbler |
| 10 | 2800m | small-bodied | *Phylloscopus chloronotus* | Lemon-rumped Warbler |
| 11 | 2800m | small-bodied | *Phylloscopus maculipennis* | Ashy-throated Warbler |
| 12 | 2800m | small-bodied | *Phylloscopus pulcher* | Buff-barred Warbler |
| 13 | 2800m | small-bodied | *Phylloscopus reguloides* | Blyth's Leaf Warbler |
| 14 | 2800m | small-bodied | *Pteruthius xanthochlorus* | Green Shrike-Babbler |
| 15 | 2800m | small-bodied | *Regulus regulus* | Goldcrest |
| 16 | 2800m | small-bodied | *Sitta himalayensis* | White-tailed Nuthatch |
| 17 | 2800m | small-bodied | *Sylviparus modestus* | Yellow-browed Tit |
